# *De novo* transformer modeling improves recovery of genetic cell types from sparse single-cell RNA sequencing

**DOI:** 10.64898/2026.08.03.742620

**Authors:** Jack M. Craig, M. Erfan Mowlaei, Sayaka Miura, Xinghua Shi, Sudhir Kumar

## Abstract

Single-cell RNA sequencing (scRNA-seq) simultaneously provides gene-expression profiles and genetic variants from individual cells, creating an opportunity to relate cellular phenotypes to their somatic evolutionary histories. However, delineating genetic types (GTs) from scRNA-seq remains difficult because most bases remain undetected in each cellular genome and are accompanied by false-positive and false-negative base calls. We evaluated some existing phylogenetic and imputation methods for GT inference using one simulated and two empirical tumor datasets. We found that the extreme sparsity of cell-variant (CV) alignments produced by scRNA-seq data prevented reliable GT recovery, particularly when multiple GTs were present. This led us to explore a transformer-based genotype imputation system (STICI), which has been shown to reduce sparsity in germline genomes. However, STICI was trained on germline sequence variation in the human population, making it inapplicable for imputing idiosyncratic somatic variation in tumor genomes. We tested the hypothesis that a STICI model trained on observed bases in a given CV alignment can predict missing bases in that dataset. We found that sample-specific STICI (ssSTICI) models predicted many missing bases with high imputation probabilities, and the resulting CV alignments placed the constituent cells of each GT closer together in the inferred cellular phylogenies. Therefore, sample-specific deep learning models show strong potential to reduce CV alignment sparsity, paving the way for integrative studies of sequence and expression evolution using scRNA-seq and similar technologies.

## Introduction

Single-cell RNA sequencing (scRNA-seq) has transformed the study of cellular heterogeneity by enabling genome-wide measurement of gene expression in individual cells from the same sample (Mereu et al., 2020; Moravec et al., 2023). Analyses of scRNA-seq expression profiles have yielded fundamental insights into cellular differentiation during development, tissue organization, and the initiation and progression of diseases such as cancer (Potter, 2018; Lim et al., 2020; Tirosh and Suva, 2024; Boxer et al., 2025). As a result, scRNA-seq has become a key technology for investigating tumor evolution and cellular diversity.

Beyond measuring gene expression, scRNA-seq reads can also infer genetic variants in expressed regions of individual cells. Consequently, every scRNA-seq experiment can provide two complementary sources of biological information: (a) gene-expression profiles that describe cells’ functional state and (b) somatic genetic variants that may help infer genetic grouping of cells. Current analyses routinely exploit the first source by clustering cells into expression-based cell types (ETs) (Hwang et al., 2018; Kiselev et al., 2019; Zhang et al., 2019; Heumos et al., 2023). However, this second source of information remains difficult to use to delineate clusters of genetically related cell groups, or genetic types (GTs). If GTs can be inferred reliably, then the joint analysis of ETs and GTs will make it possible to determine whether similar expression states arise through common ancestry, transitions during cellular differentiation, or convergent evolution among genetically distinct cell populations (Patel et al., 2014; Chung et al., 2017; Puram et al., 2017; Kim et al., 2020; Huzar et al., 2022). Such joint analyses will, thus, provide a direct framework for understanding how genetic evolution gives rise to somatic cellular phenotypes during normal development and disease progression.

Current studies often infer GTs from large-scale genomic alterations, particularly copy-number alterations (CNAs), because scRNA-seq detects these events more reliably than individual nucleotide variants (Hadi et al., 2020; Tarabichi et al., 2021). Consequently, CNA-defined cell clusters have become a practical surrogate for GTs in studies of tumor evolution (Gao et al., 2021). Although highly informative and increasingly refined (Gayoso et al., 2022; Jun et al., 2023), CNAs are relatively infrequent macro-mutational events and therefore provide only coarse genetic resolution. By contrast, single-nucleotide variants (SNVs) are far more numerous and provide fine resolution for evolutionary inference (Beroukhim et al., 2010; Zack et al., 2013; ICGC/TCGA, 2020). If accurately recovered from scRNA-seq datasets, delineation of GTs using SNVs would offer high-resolution studies of genetic and expression evolution of tumors.

For SNV-based GT inference, we can assemble the SNVs detected in scRNA-seq data into a cell-variant (CV) alignment and then apply molecular phylogenetic methods to infer evolutionary clusters corresponding to GTs. CV alignments are sparse (**Fig. 1**) because scRNA-seq techniques can detect only a small fraction of the transcripts expressed in each cell, and only a subset of these transcripts contain informative variants (Ziegenhain et al., 2017; Mereu et al., 2020; Chen et al., 2021). Consequently, CV alignments have abundant missing observations for cell-site combinations, even in high-quality datasets (Sparta et al., 2024). Sequencing and variant-calling procedures also introduce false-positive and false-negative base calls (Svensson et al., 2017; Schnepp et al., 2019; Lu et al., 2021). As a result, current methods struggle to reliably recover cellular genetic relationships and GTs from raw CV alignments (Liu et al., 2023; Moravec et al., 2023; Dou et al., 2024).

**Figure 1:**
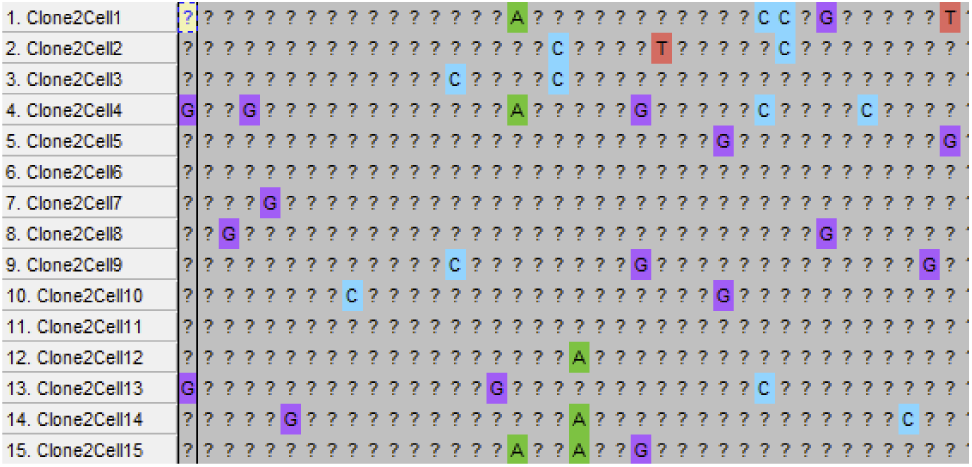
A representative multiple sequence CV alignment derived from scRNA-seq data. An scRNA-seq experiment typically produces highly sparse CV alignments where some cells have base calls at a site (shaded boxes), whereas most positions are unobserved (“?”).

Several computational methods address missing data and sequencing errors in sparse CV alignments in distinct ways. BEAM uses phylogenetic relationships among evolutionarily proximate cells to iteratively improve base assignments (Miura et al., 2018), PhylinSic employs probabilistic smoothing to infer missing information in scRNA-seq datasets (Liu et al., 2023), and CellPhy jointly models genotype evolution and sequencing error during phylogenetic inference (Kozlov et al., 2022). Nonetheless, recovering the evolutionary signal needed for successful base prediction becomes challenging when CV alignments are very sparse, limiting accurate delineation of GTs (see *Results*).

Recent advances in machine learning suggest a fundamentally different strategy for recovering genetic information from sparse, noisy datasets (Arisdakessian et al., 2019; Chen and Shi, 2019; Eraslan et al., 2019; Badsha et al., 2020; Dias et al., 2022). Transformer models learn complex dependencies across sequence positions and accurately predict missing genomic variants from incomplete observations (Devlin et al., 2018; Mowlaei et al., 2025). Their success in imputing variants in regions with high linkage disequilibrium (LD) with extraordinary accuracy (Mowlaei et al., 2025) suggests they may also recover missing sample-specific somatic variants from sparse CV alignments, because somatic mutations in a cellular lineage are inherited through successive mitotic cell divisions without recombination (i.e., perfect LD). Consequently, related cells preserve characteristic patterns of variant co-occurrence that reflect their shared ancestry. Even when only a small fraction of variant positions is directly observed, these lineage-dependent patterns may provide sufficient information for transformer models to infer many missing bases within a CV alignment.

We initially considered applying STICI, a transformer architecture which has shown promise in genotype imputation using global population reference panels (Mowlaei et al., 2025). However, STICI was trained on germline variants collected from many individuals across diverse human populations, whereas scRNA-seq datasets contain sample-specific somatic variants that are largely unique to an individual tumor and, therefore, scarce in existing reference panels. This distinction prompted us to investigate whether we could train a STICI model *de novo* for a given CV alignment generated from a sample of tumor cells from one individual. The sample-specific STICI (ssSTICI) would learn observed patterns of somatic variant co-occurrence directly from the given sample (CV alignment), which would then inform imputation of unobserved bases within that same alignment. An ssSTICI model trained on one CV alignment may not apply to another CV alignment from a different tumor sample or individual because the cells may not share the same phylogeny and may have different SNVs. So, an ssSTICI model is trained on observed bases from the same CV alignment it uses to predict missing bases. Here, we first evaluate existing methods’ ability to recover GTs from highly sparse scRNA-seq CV alignments using one simulated dataset and two empirical tumor datasets. We then describe ssSTICI modeling for individual datasets and evaluate whether the predicted missing bases can improve GT delineation.

## Results

### Existing methods fail to recover GTs from sparse CV alignments

We asked whether existing approaches can reliably delineate GTs from the highly sparse CV alignments generated by scRNA-seq experiments. To address this question, we simulated somatic evolution along a clonal phylogeny containing five genetically distinct GTs, each represented by 1,000 genetically identical cells, with 500 variant positions. We modified the CV alignment to reproduce scRNA-seq data characteristics by replacing 95% of bases with the missing symbol (?) and imposing false-positive and false-negative sequencing errors typical of scRNA-seq profiles (see *Methods*).

To recover the known clonal structure from this simulated dataset, we inferred a molecular phylogeny using FastTree because of the large number of cells (Price et al., 2010). Rather than forming five distinct clusters, cells from individual GTs were scattered throughout the phylogeny (**Fig. 2** “Raw” trees). To quantify cell clustering according to their GTs, we used two complementary measures: the neighborliness index (*NI*) and the distance ratio (*DR*). *NI* is the fraction of cells whose nearest topological neighbor in the phylogeny belongs to the same GT. An *NI* of 1.0 indicates that every cell is adjacent to a member of the same GT, whereas lower values indicate increasingly poor local clustering. *DR* is the average pairwise patristic distance between cells from different GTs in the inferred phylogeny divided by the average pairwise distance among cells within the same GT. Higher *DR* values indicate stronger GT separation and tighter clustering within each GT (see *Methods*).

**Figure 2:**
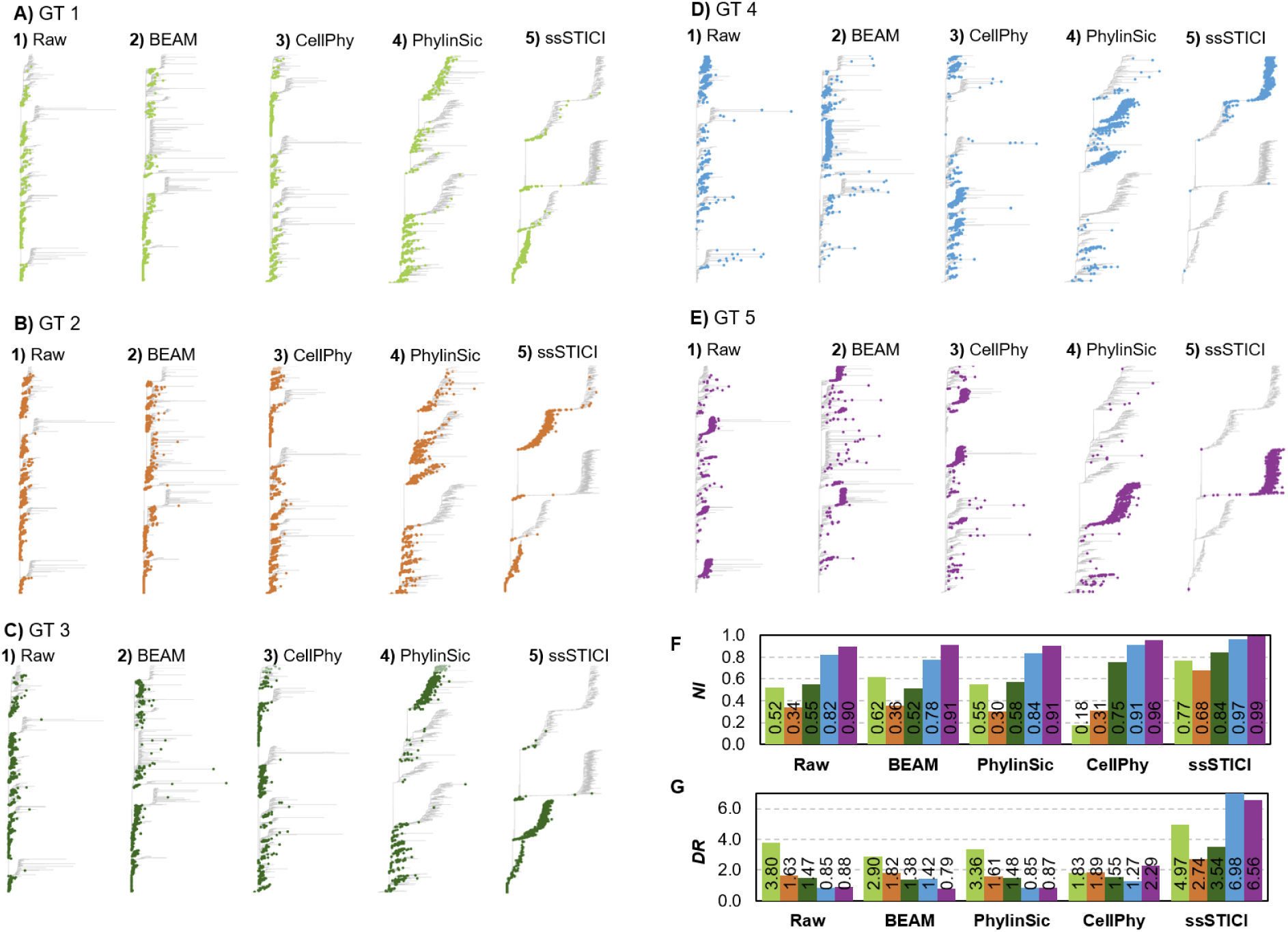
Recovery of five true GTs in the simulated dataset. For each GT (**A-E**, color-coded), the corresponding cells (1,000 per GT) are highlighted on the phylogeny inferred **1**) from the CV alignment with sparsity and error added; **2**) after BEAM analysis; **3**) after CellPhy analysis; **4**) after PhylinSic analysis; and **5**) after ssSTICI analysis. **F)** *NI* scores: Raw average = 0.63; BEAM average = 0.64; PhylinSic average: 0.63; CellPhy average = 0.62; ssSTICI average = 0.85. **G)** *DR* scores: Raw average = 1.73; BEAM average = 1.66; PhylinSic average: 1.63; CellPhy average = 1.77; ssSTICI average = 4.96. In the case of all GTs, the true cellular sequences are identical, so true within-GT branch lengths are zero.

In the phylogeny inferred from the raw CV alignment (raw phylogeny), *NI* ranged from 0.34 to 0.90, averaging 0.63 across GTs, which is well below the target of 1.0. *DR* values were also low and fell below 1.0 for some GTs, indicating that, on average, cells from different GTs were placed closer than members of the same GT in the raw phylogeny. Thus, the raw phylogeny poorly recovered the known GTs, consistent with reports of challenges in empirical scRNA-seq datasets (Moravec et al., 2023).

We next evaluated complementary existing methods, including BEAM (Miura et al., 2018), PhylinSic (Liu et al., 2023), and CellPhy (Kozlov et al., 2022). BEAM produced few confident imputations at the default Bayesian posterior probability (*BPP*) threshold of 0.7, because BEAM relies on phylogenetic signal in the starting CV alignment, which was rather limited as seen in the Raw phylogeny (**Fig. 2**). This is often tractable for datasets obtained using single-cell DNA sequencing, which have less than 50% sparsity (Miura et al., 2018), but CV alignments from scRNA-seq typically exceed 90% sparsity. After BEAM analysis, average *NI* for GTs in the inferred phylogeny changed by only 0.01, and *DR* did not improve, as the cells were dispersed as much as in the raw phylogeny (**Fig. 2**).

After PhylinSic processing, average *NI* remained 0.63, while average *DR* changed from 1.73 to 1.63 **(Fig. 2)**. This is because clustering improved slightly for GT1 and GT4, but GT5 cells appeared more dispersed. CellPhy noticeably improved some GTs (**Fig. 2**). GT5 (purple) became a remarkably cohesive cluster, and GT4 (blue) improved substantially. However, GT1 and GT2 worsened. Overall, the average *NI* was lower than in the raw phylogeny, while the average *DR* improved (**Fig. 2**). Thus, none of the tested methods consistently recovered the simulated tumor’s known GTs for practical data analysis.

### ssSTICI recovers information from sparse CV alignments

We trained an ssSTICI model for the CV alignment from the simulated dataset (see *Methods*). During its training, we randomly masked 80% of the observed bases, following Mowlaei et al. (Mowlaei et al., 2025). Because the simulated data retained only 5% of the bases in the CV alignment, different combinations of 4% masked bases and 1% unmasked bases were used in training iterations. During this training, the CV alignment had 95% missing bases (which were held out). In these iterations, the model strived to correctly predict the masked (4%) bases using the unmasked observations (1%). Training an ssSTICI model for this dataset required 3.3 hours and 0.6 GB of VRAM on an NVIDIA H100 GPU.

We then used the trained ssSTICI model to impute the missing data (95% missing bases), which required less than 1 minute on the same machine. It predicted softmax probabilities for each base at every position in every sequence, which we refer to as transformer base probabilities (*TBPs*). Base-prediction accuracy rose monotonically with higher *TBP* thresholds, but the number of imputed bases declined, especially above a *TBP* threshold of 0.99 (**Table 1**). *NI* and *DR* of phylogenies inferred using the resulting CV alignments rose slightly with the *TBP* threshold up to 0.99, past which point *NI* decreases dramatically for most GTs and *DR* becomes erratic (**Fig. 3**). Consequently, we imposed a *TBP* filtering threshold of 0.99 on all datasets for GT inference to balance GT recovery and base call prediction accuracy.

**Figure 3.**
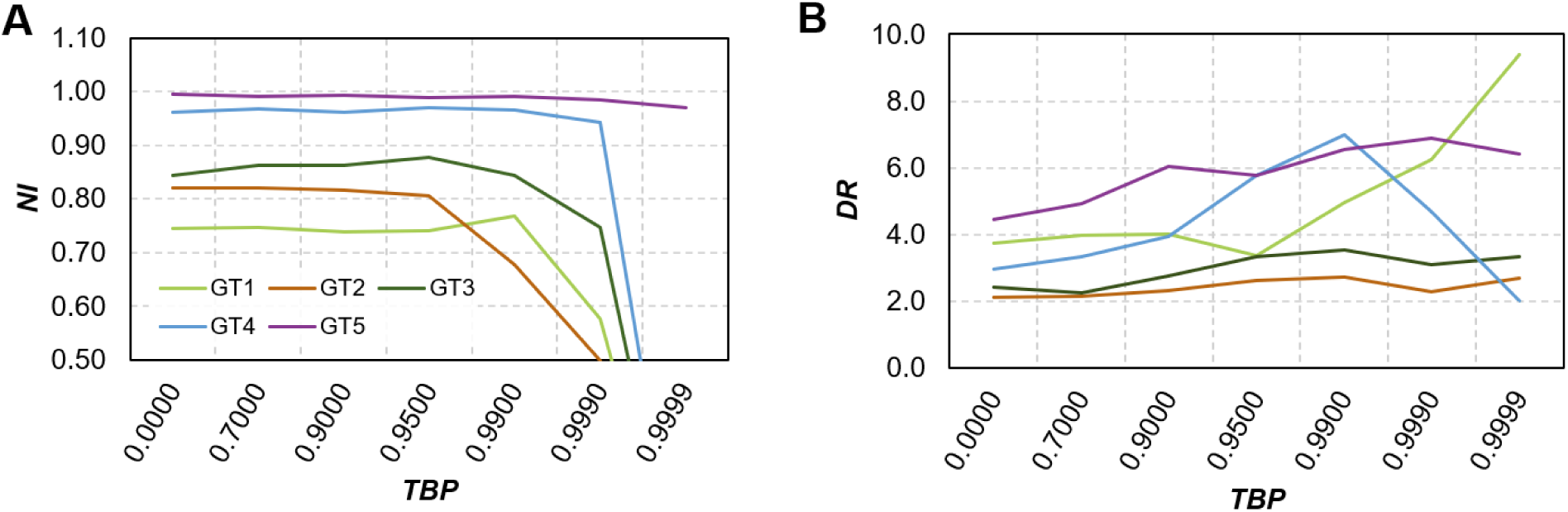
Sensitivity of *NI* and *DR* metrics to transformer base probability (*TBP*). We tested a range of *TBP* filtering thresholds ranging from 0 (the highest probability base call was always accepted at every position) to 0.9999 (all predictions with a *TBP* < 0.9999 were rejected). We observe that both *NI* (**A**) and *DR* (**B**) improve slightly as the *TBP* threshold increases up to 0.99, after which *NI* drops dramatically for most GTs, and *DR* becomes erratic, increasing for some and decreasing for others. This behavior supports the *TBP* filtering threshold of 0.99 we imposed on all datasets for GT inference.

**Table 1:**
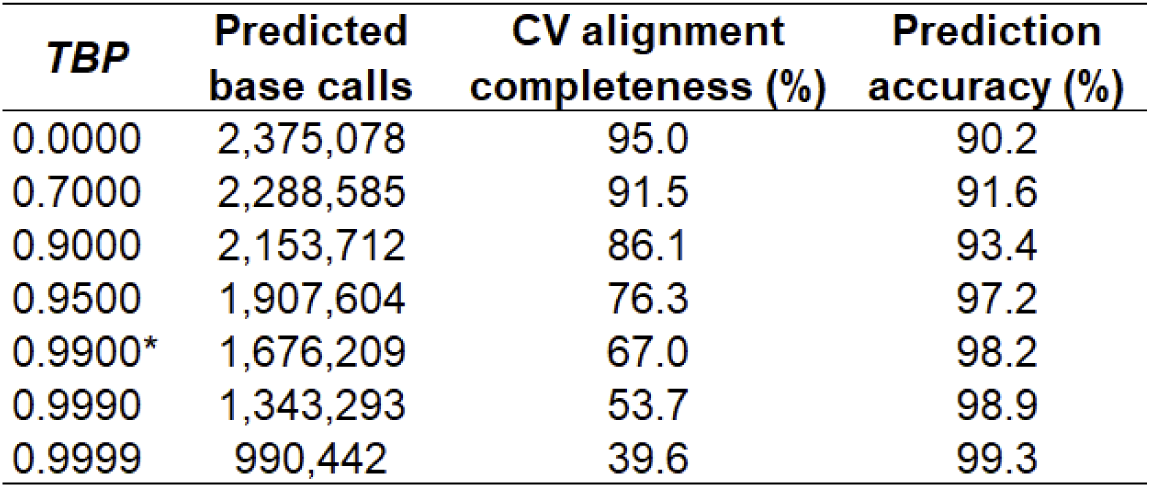
Effect of transformer base probability (*TBP*) filtering on quantity and accuracy of predicted base calls in the simulated dataset. Prediction accuracy increases with *TBP* filter stringency, but CV alignment completeness declines sharply after a *TBP* of 0.99. The “*” indicates the *TBP* filtering threshold used for all tree-building and GT recovery tests. Base calls observed in training (5%) were excluded, so maximum CV alignment completeness is 95%.

| *TBP* | Predicted<br>base calls | CV alignment<br>completeness (%) | Prediction<br>accuracy (%) |
| --- | --- | --- | --- |
| 0.0000 | 2,375,078 | 95.0 | 90.2 |
| 0.7000 | 2,288,585 | 91.5 | 91.6 |
| 0.9000 | 2,153,712 | 86.1 | 93.4 |
| 0.9500 | 1,907,604 | 76.3 | 97.2 |
| 0.9900* | 1,676,209 | 67.0 | 98.2 |
| 0.9990 | 1,343,293 | 53.7 | 98.9 |
| 0.9999 | 990,442 | 39.6 | 99.3 |

### Recovery of missing bases substantially improves genetic cell typing

#### Simulated dataset

Using ssSTICI for the sparse CV alignment produced from the simulated dataset, base-prediction accuracy for 1,676,209 predicted bases was 98.2% at a *TBP* threshold of 0.99. However, using ssSTICI to predict bases at positions in the original CV alignment used to train the model yielded lower accuracy (93.0%). This is because the model ended up learning to recover some of the erroneous 21% of the 5% bases used for training, as they were sometimes assigned a high *TBP* during the inference pass (**Fig. 4**). Since this error was randomly distributed in the training data, it did not leave any imprints on the base dependency patterns learned by the ssSTICI model, and allowed it to correctly impute a vast majority of the 95% missing bases in the CV alignment. So, in all future analyses, we excluded the observed base calls used for training from the CV alignments used to delineate GTs.

**Figure 4.**
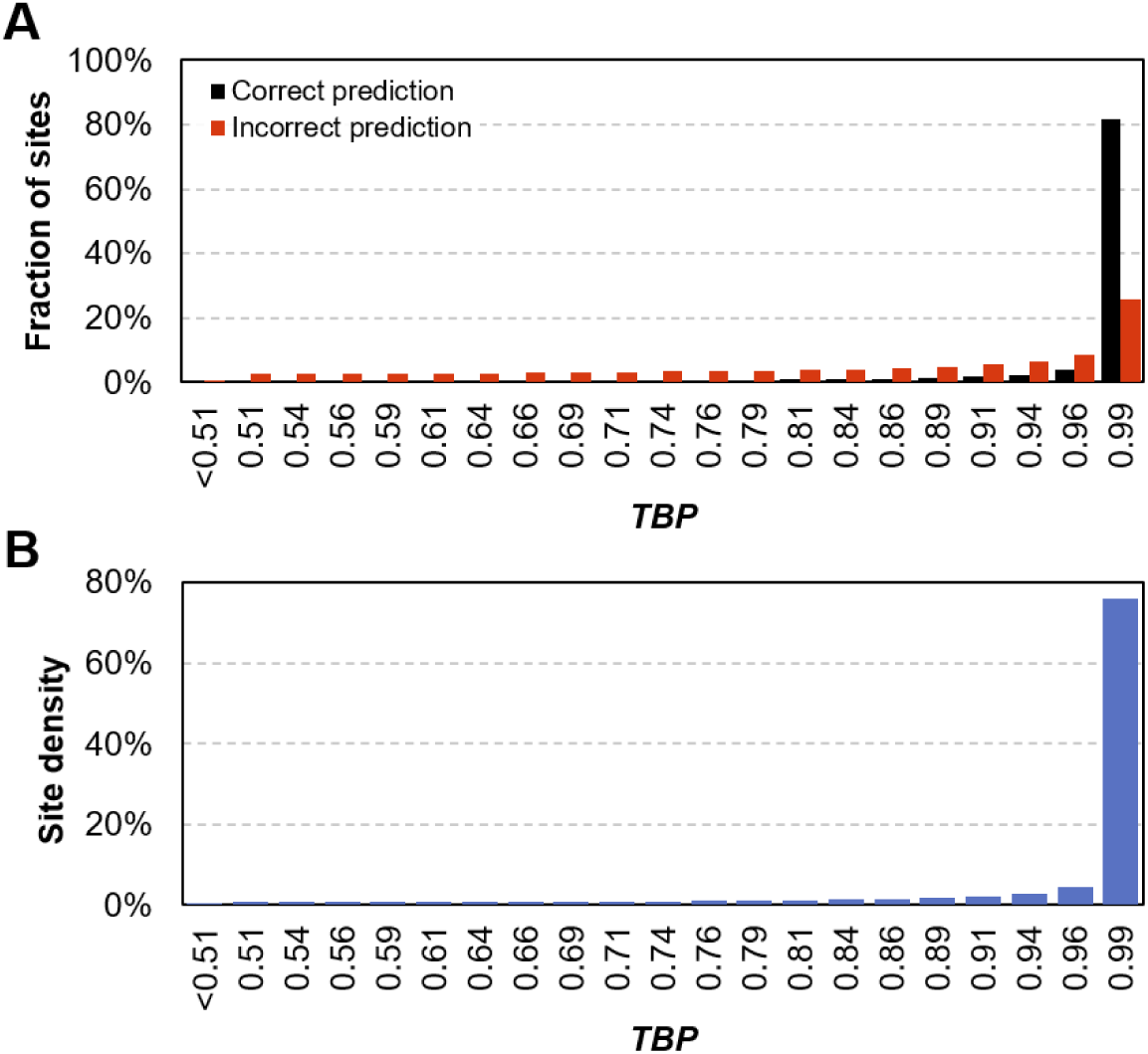
Distribution of transformer base probabilities (*TBPs*). **A)** Distributions of *TBP* scores for positions with correct base predictions (black) and incorrect predictions (red). Incorrect predictions are far more likely at lower *TBPs*. **B)** Density of sites by *TBP* of the prediction. The abundance of sites with *TBPs* >0.99 and the substantially greater accuracy of base calls with *TBPs* >0.99 support the use of this threshold for filtering prior to GT inference.

The CV alignment produced using ssSTICI greatly improved clustering for each GT (**Fig. 2**). The *NI* of GT1, GT2, and GT3 increased by over 0.20 each, from 0.52 to 0.77, 0.34 to 0.68, and 0.55 to 0.84, respectively. The *NIs* of GT4 and GT5, which were initially the most well-defined, improved less, from 0.82 to 0.97 and 0.90 to 0.99, respectively (**Fig. 2**). The average *NI* increased from 0.63 to 0.85, with each GT’s *NI* exceeding the average *NI* of every other method.

The *DR* values improved more dramatically: GT4 increased from 0.85 to 6.98, and GT5 from 0.88 to 6.56. In both cases, the distance ratio inverted completely due to long internal stem branches in the inferred phylogeny. GT1, GT2, and GT3 also improved, from 3.80 to 4.97, 1.63 to 2.74, and 1.47 to 3.54, respectively (**Fig. 2**). The average *DR* increased to 4.96, double that of any other method. Thus, the use of ssSTICI substantially improved recovery of simulated GTs compared with existing methods.

We further tested a random permutation of CV alignment positions and cells by randomizing the column and row order and used ssSTICI to predict missing base calls (see *Supplementary Materials*). We observed only marginal changes in *NI* and *DR* (<5%), suggesting that ssSTICI was robust to changing the alignment order in this case.

To determine the sensitivity of ssSTICI to unobserved positions in the input alignment, we also conducted a threefold cross validation in which three new ssSTICI models were trained, each with all base calls masked at a randomly-selected third of the sites in the input (see *Supplementary Materials*). We observed no substantial decline in prediction accuracy at those sites for any model, and GT inference was not meaningfully impacted either (**Table S1**).

#### Ovarian cancer dataset

We analyzed an ovarian cancer (OC) dataset containing 2,970 SNVs in 1,827 cells of six tumors sampled from different anatomical sites, including primary and metastatic tumors (Vázquez-García et al., 2022). CopyKAT (Gao et al., 2021) detected two major GTs based on CNAs, which we used as an *ad hoc* ground truth for validation in the absence of simulated GTs. The initial CV alignment was missing 86.0% of the bases, and its raw phylogeny had an average *NI* of 0.84 and *DR* of 1.11. Cells from two CNA-defined GTs did not cluster cohesively in the phylogeny inferred from the initial CV alignment (**Fig. 5**).

**Figure 5:**
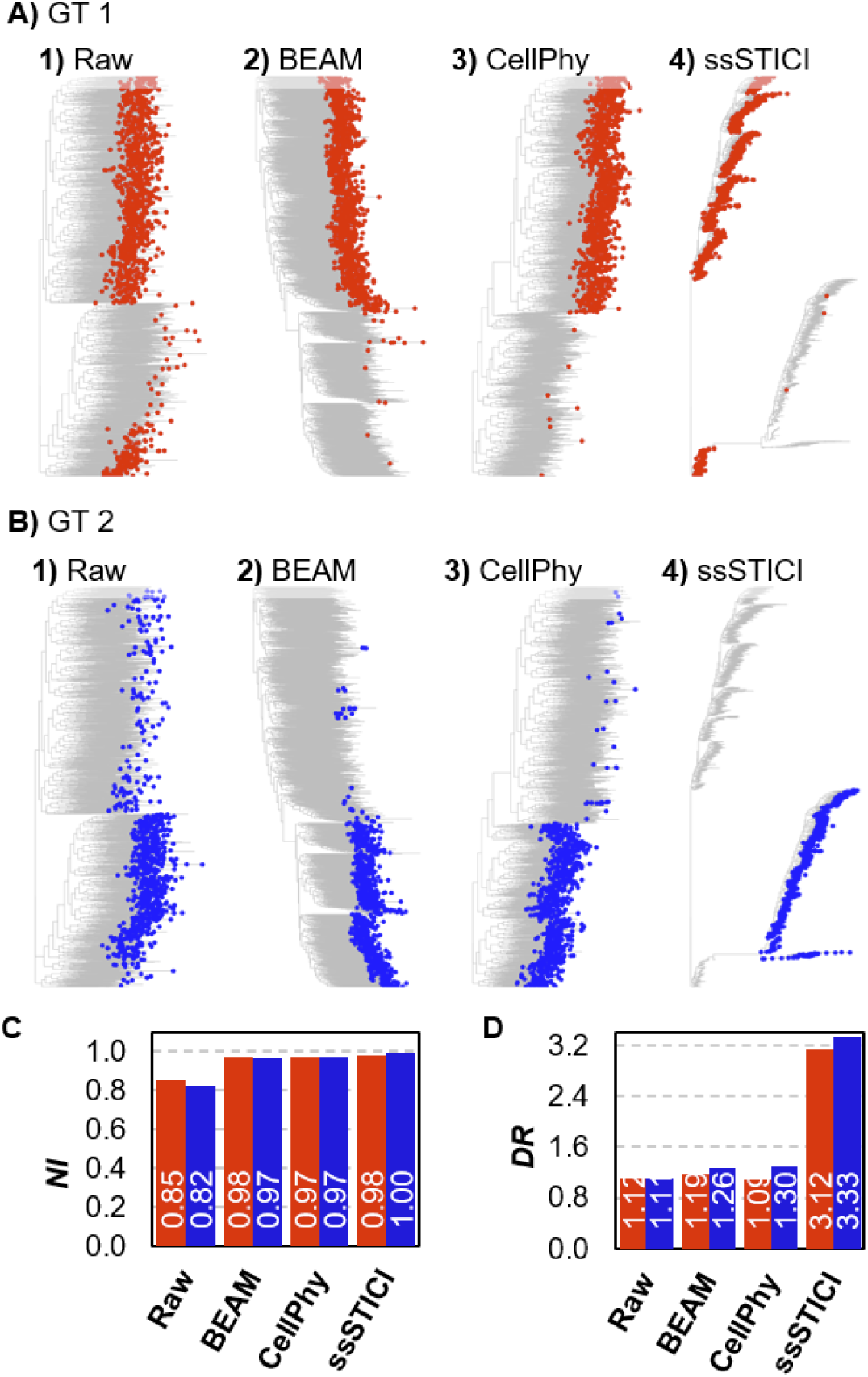
Recovery of two CNA-defined GTs in the empirical OC dataset. For each GT (**A** and **B**, color-coded), the corresponding cells (GT1 = 1,058 cells; GT2 = 769 cells) are highlighted on the phylogeny inferred **1**) from the raw scRNA-seq dataset; **2**) after BEAM analysis; **3**) after CellPhy analysis; **4**) after ssSTICI analysis. **C)** *NI* scores: Raw average = 0.84; BEAM average = 0.97; CellPhy average = 0.97; ssSTICI average = 0.99. **D)** *DR* scores: Raw average = 1.11; BEAM average = 1.23; CellPhy average = 1.17; ssSTICI average = 3.23.

BEAM and CellPhy recovered the two major CNA-defined GTs reasonably well (**Fig. 5**). Only a few cells were misclassified, resulting in average *NI values* of 0.97 and *DR* values greater than 1 for both. PhylinSic could not be run on a dataset of this size. Thus, existing approaches could perform well when only two major GTs must be distinguished,

Training the ssSTICI model for the OC dataset required about 1 hour and 4.9 GB of VRAM on an NVIDIA H100 GPU. It involved many iterations of masking 80% of the 14% observed bases. At the *TBP* threshold of 0.99, the ssSTICI model imputed 1,732,685 missing bases, reducing sparsity from 86.0% to 54.4%. The resulting CV alignment yielded greatly improved separation of the two CNA-defined GTs (**Fig. 5**). Average *NI* increased to 0.99, while *DR* increased from 1.11 to 3.23, revealing much higher success in clustering cells of the two GTs (**Fig. 5**). The threefold increase in *DR* is readily apparent from the tree itself, as no existing method could clearly discriminate between clones, whereas CV derived using ssSTICI predictions places GT2 on a long stem branch of characteristic SNVs. Likewise, existing methods all yielded long pendant edges composed of (likely spurious) unique SNVs and subtending individual cells, whereas ssSTICI minimized these, further improving *DR* (**Fig. 5**).

Here, the permutation test reduced performance relative to the unaltered CV alignment (see *Supplementary Materials*). *NI* was largely unaffected, but *DR* dropped by 52.4% on average, largely due to shortening of the internal branches which discriminate between GT. This suggests ssSTICI may have some sensitivity to site order for longer CV alignments.

#### Triple-negative breast cancer dataset

We also analyzed a triple-negative breast cancer (TNBC) dataset containing 3,425 cells, 19,576 sites, and six CNA-defined GTs (Gao et al., 2021). The CV alignment was 92% sparse, producing a raw phylogeny with an average *NI* of 0.27 and *DR* of 1.06 (**Fig. 6**). BEAM improved *NI* for several GTs, especially GT2, which increased from 0.30 to 0.79, GT3, which increased from 0.03 to 0.23, and GT6, which increased from 0.26 to 0.82. *NI* values for GT1, GT4, and GT5 increased by 0.25–0.50 each (**Fig. 6**). For three clones, *DR* improved from essentially random to slightly clustered: GT2 increased from 0.99 to 1.55, GT3 from 1.01 to 1.23, and GT4 from 1.02 to 1.41. In GT1, GT5, and GT6, *DR* essentially did not change (**Fig. 6**). PhylinSic could not scale to this dataset, and CellPhy analyses did not complete after several weeks. Therefore, methods that performed reasonably well on the two-GT OC dataset were much less effective for this multi-GT problem (BEAM) or not scalable to this dataset size (CellPhy and PhylinSic).

**Figure 6:**
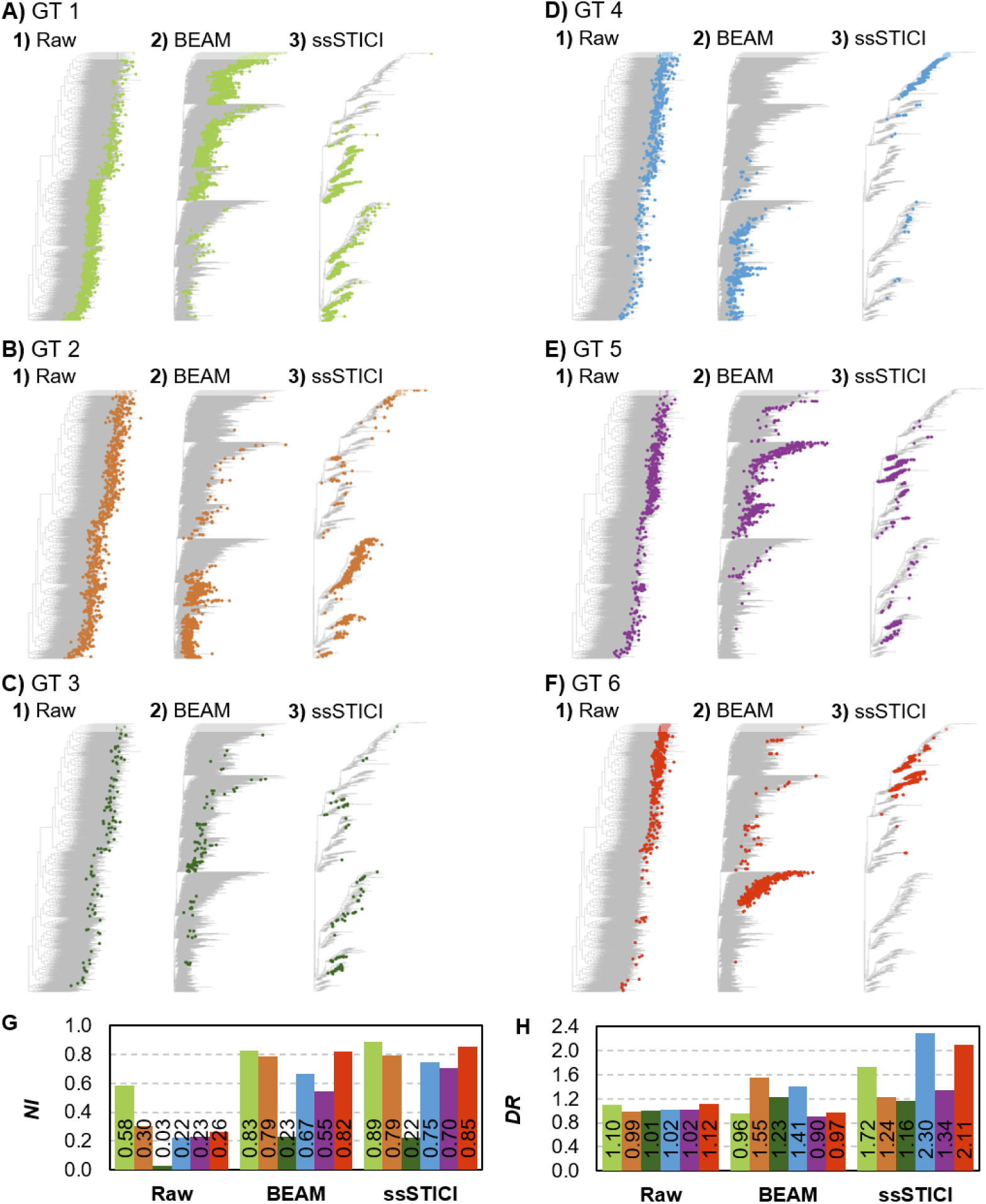
Recovery of six CNA-defined GTs in the empirical triple-negative breast cancer dataset. For each GT (**A-F**, color-coded), the corresponding cells (GT1 = 1,309 cells; GT2 = 750 cells; GT3 = 104 cells; GT4 = 400 cells; GT5 = 449 cells; GT6 = 413 cells) are highlighted on the phylogeny inferred **1**) from the raw scRNA-seq dataset; **2**) after BEAM analysis; **3**) after ssSTICI analysis. **G)** *NI* scores: Raw average = 0.27; BEAM average = 0.65; ssSTICI average = 0.70. **H)** *DR* scores: Raw average = 1.06; BEAM average = 1.17; ssSTICI average = 1.64.

Training an ssSTICI model on this larger dataset required about 10 hours and 38.0 GB of VRAM on an NVIDIA H100 GPU. At a *TBP* threshold of 0.99, the ssSTICI model predicted 20,605,000 missing bases, reducing CV alignment sparsity to 61.9%. The resulting CV alignment improved *NI* substantially more than BEAM for GT1 (increased to 0.89), GT4 (0.75), and GT5 (0.70), while GT2, GT3, and GT6 saw essentially the same improvement in *NI* with ssSTICI as with BEAM. Improvements were greater for *DR:* ssSTICI raised GT1 to 1.72, GT5 to 1.34, and GT6 to 2.11, whereas BEAM reduced *DR* for these GTs below 1. Relative to BEAM, *DR* changed from 1.55 to 1.24 for GT2 and from 1.23 to 1.16 for GT3, while GT4 improved from 1.41 to 2.30. The overall average *DR* was 1.64 (**Fig. 6**). As in the OC dataset, these large improvements are attributable to ssSTICI yielding longer internal branches to discriminate between GTs, paired with shorter pendant branches for each cell.

Interestingly, cells within the same CNA-defined GT clustered into small, cohesive groups (**Fig. 6**). For example, GT1 and GT6 appear to have at least two nested sub-GTs, based on the clustering of associated cells. These patterns suggest SNV-defined substructure within the broader CNA-defined GTs and were not apparent in the raw or BEAM phylogenies. Because independent SNV-based clonal assignments are unavailable, these groups should be considered as possibilities rather than confirmed biological lineages.

The permutation test for this dataset produced mixed results (see *Supplementary Materials*). *NI* increased by 6.0% while *DR* decreased by 18.3%.

## Discussion

The increasing availability of scRNA-seq data offers substantial promise for deciphering GTs and analyzing convergence and divergence of gene expression within and between GTs. Our results show that ssSTICI modeling can outperform traditional statistical approaches in delineating GTs, because it can impute a substantial fraction of missing bases with high softmax probabilities. This is because somatic cells divide by mitosis without recombination, so mutations are inherited together across generations of cells, preserving characteristic patterns of base co-occurrence. Successive mutations also generate nested combinations of variants as cells diversify. A transformer (STICI) learns base contexts in the specific CV alignment, which can then be applied to impute a majority of missing data with high confidence, compared with classical methods that do not account for dependence among sites and treat each position in the CV alignment as independent information during statistical analysis; see also Kumar et al. (Kumar et al., 2026). As expected, ssSTICI performance was mostly robust to reordering columns in the CV alignment because it shuffles base locations rather than their dependencies (see *Supplementary Results*). Furthermore, peak VRAM during training remained under 40 GB even for the largest CV alignment (**Fig. S2**). This means that many scRNA-seq alignments should be readily analyzable without advanced hardware, even on well-equipped desktop workstations.

However, ssSTICI did not correct all erroneous bases in the starting CV alignment. The model treated observed errors as correct targets and could memorize them (**Fig. 4**), which is a known issue in machine learning with noisy input data (Pompanon et al., 2005; Harutyunyan et al., 2020). However, this memorization was limited to the observed bases in the input CV alignment because we introduced sequencing errors randomly. This means that while we could not correct most errors in the initial base calls, we could impute missing bases well and use them for GT analysis. This error “smoothing” is a unique benefit of the transformer architecture, as other de-noising tools have also shown (Chi and Deng, 2020; Xia et al., 2021; Ng et al., 2023).

Three limitations of our approach are worth mentioning. First, ssSTICI models performed substantially better than classical methods when the clonal structure was more complex. BEAM and CellPhy recovered the two major CNA-defined GTs reasonably well in the OC dataset. But BEAM was much less successful in separating six GTs in the TNBC dataset, and CellPhy and PhylinSic were computationally infeasible at that scale. Thus, the difficulty is exacerbated for classical methods when sparse data need to resolve many genetically related lineages. Second, while we treated CNA-defined GTs as a useful independent benchmark in tests of empirical datasets, they are not an exact ground truth for SNV-defined GTs. CNA designations and the cell clusterings they produce are subject to known errors (Gao et al., 2021; Song et al., 2025). Furthermore, CNA-defined GTs offer a coarser view of evolution than more common SNVs, a distinction especially relevant in the TNBC analysis, where ssSTICI modeling suggests subclones separated by substantially long internal branches. Third, unobserved base calls in the simulated data were induced at random (missing completely at random, MCAR), and in empirical data, missingness may be due to real low expression (missing not at random, MNAR). Since there is no ground truth for the empirical data, the tests we do here do not directly establish the robustness of ssSTICI to expression-dependent dropouts. Future tests of missing not at random (MNAR) dropouts will be needed to determine this.

Nevertheless, our results establish the potential of transformer-based GT inference from highly sparse scRNA-seq data. Future methods, particularly advanced transformer architectures, will need to address lingering issues, such as distinguishing uncertainty from missing observations versus uncertainty from sequencing and variant-calling errors. Information such as read depth, allele balance, base quality, and mapping quality may help determine how strongly an observed call should contribute to training and inference.

Methods will also need to be tested across datasets containing different numbers and arrangements of GTs, because the contrast between the two-GT and six-GT analyses indicates that performance can decline as lineage complexity increases. When transformer architectures mature sufficiently to address these issues, we expect GT inference to be highly reliable even from sparse scRNA-seq datasets.

## Methods

### Simulated data

We simulated a sparse, error-prone CV alignment by adapting an existing simulated dataset from a previous study that benchmarked methods for inferring cellular phylogeny (Miura et al., 2023). This dataset consisted of five GTs, each with 1,000 cells, and each cell was characterized by 500 SNVs. The original simulation incorporated a 1% false-positive rate and a 20% false-negative rate (Miura et al., 2023). To mimic the sparsity observed in empirical scRNA-seq data, we randomly replaced bases with missing values until overall sparsity reached 95%. Because the original dataset encoded only the presence or absence of mutations, we randomly assigned nucleotide states to convert it into sequence data.

### Empirical data

We obtained empirical scRNA-seq data from prior studies of metastatic OC tumor cells (Vázquez-García et al., 2022) and triple-negative breast cancer (TNBC) tumor cells (Gao et al., 2021). For the OC patient, the original study sequenced six tumors sampled from different anatomical sites, including primary and metastatic tumors, resulting in six biological samples from the same patient. For the TNBC patient, the original study sequenced a single tumor sample, corresponding to one sequencing library. We analyzed the OC and breast cancer datasets separately and did not pool them.

For each sample, we processed raw sequencing reads (FASTQ files) into gene-expression matrices using the Cell Ranger Count pipeline from 10x Genomics (Genomics, 2019). We also generated position-sorted read alignments (BAM files) for downstream SNV calling. In both cases, these datasets represent tumor cell populations sequenced in a single experiment from the same individual. For gene-expression analysis, we removed ambient RNA contamination, doublets or multiplets, and low-quality cells with elevated mitochondrial RNA content using SoupX (Young and Behjati, 2020), DoubletFinder (McGinnis et al., 2019), and miQC (Hippen et al., 2021), respectively. We performed cell-type annotations using SingleR (Aran et al., 2019), with annotations from the original study for the OC dataset and from the Human Breast Cell Atlas for the TNBC dataset (Kumar et al., 2023).

For the OC dataset, we integrated samples in Seurat to identify SNV positions and CNAs. CNAs were inferred using CopyKAT (Gao et al., 2021). Ploidy status was estimated for each cell, and only cells inferred by CopyKAT to be aneuploid were retained for CNA-defined GT annotation.

For SNV calling, we used OC cells in the OC dataset and luminal epithelial and basal mammary cells in the TNBC dataset. We identified SNVs using cellSNP-lite (Huang and Huang, 2021) with default settings, including a minimum minor-allele frequency of 0.1 and minimum read depth of 100. SNVs with frequencies greater than 0.5% were retained for phylogenetic inference. After processing, the OC dataset contained 1,827 cells and 2,970 SNVs, whereas the TNBC dataset contained 3,425 cells and 19,576 SNVs.

### Existing Methods

We analyzed simulated and empirical datasets with BEAM (Miura et al., 2018) using the BEAMfast.py implementation and default parameters. We used PhylinSic (Liu et al., 2023) with default parameters, and CellPhy (Kozlov et al., 2022) with the “fast” method for the simulated dataset, as the “full” and “search” methods were computationally infeasible. We could run all three methods on the simulated data. We could run CellPhy on the OC dataset, but we did not complete PhylinSic or CellPhy on the TNBC data because of the dataset size.

### Training and applying ssSTICI

The STICI architecture was originally designed for germline genomic imputation (Mowlaei et al., 2025), which we adapted for imputing missing bases in the input CV alignment. That is, we kept the model architecture intact, including arbitrary column-order 1D convolutions, but adjusted the data pre-processing (e.g., tokenization), post-processing, and loss functions. Out of the three loss functions used in STICI, we removed the Mach-Rsq loss function because it was designed to improve genotype-dosage prediction and was not relevant to scRNA-seq base prediction (Mowlaei et al., 2025). Consequently, ssSTICI uses categorical cross-entropy and Kullback-Leibler divergence loss functions and ignores missing sites during training. We trained the original STICI model on a complete reference panel of genome sequences from the 1000 Genomes Project. For CV alignments derived from scRNA-sequencing, we trained STICI separately for each CV alignment, i.e., we built a sample-specific model, ssSTICI. During training, 80% of the observed base calls are masked, and the remaining 20% are used by the model to predict them in iterative epochs. This means that for an alignment with only 5% observed sites, predictions of 4% of the total sites are made based on just 1% of the sites. Importantly, sites are masked randomly, and the final predictions are robust to the pattern of masked sites (see *Supplementary Results*).

**Supplementary Fig. S1** shows the full confusion matrix of base-by-base prediction accuracy against true base identities in the simulated dataset pre-and post-ssSTICI. Overall, prediction accuracy was 94.5% across all sites in the simulated dataset, where the true reference is known, but it cannot be calculated for empirical datasets. Training accuracy, calculated as the percentage of bases masked during training which the model can correctly predict by the final training epoch, was 80.4% for the OC dataset and 77.2% for the TNBC dataset. After training, ssSTICI produced a distribution of transformer base probabilities (*TBPs*) for every site in each sequence, allowing us to predict missing bases at the given TBP threshold.

We deposited the model source code, processed alignment files, and *NI*/*DR* scoring scripts used for training, prediction, and evaluation at https://github.com/ErfanMowlaei/ssSTICI.

### Assessment of clonal grouping

We used FastTree to analyze simulated, empirical, and ssSTICI-completed CV alignments because it can handle datasets with thousands of cells (Price et al., 2010). We used a GTR model (-gtr) for full-base-resolution CV alignments in the OC and simulated datasets and the standard nucleotide option (-nt) for the binary TNBC CV alignment.

We quantified clonal grouping using two metrics. For the simulated dataset, we knew the true clonal identities. For empirical datasets, we used CNA-defined GT assignments inferred by CopyKAT as an external benchmark (Gao et al., 2021). The neighborliness index (*NI*) is the fraction of cells assigned to a given GT whose nearest topological neighbor is also assigned to that GT. When the nearest topological neighbor is a clade containing multiple cells, the clonal assignment for that clade is the most common identity among its members, with ties broken at random. *NI* ranges from 0.0 to 1.0, with higher values indicating stronger local clustering of cells from the same GT. The distance ratio (*DR*) is the average pairwise patristic distance between cells from different GTs divided by the average pairwise distance among cells from the same GT. Greater distances between GTs and shorter distances within GTs therefore produce higher *DR* values, indicating stronger recovery of clonal structure.

