## Supplementary materials and figures for "*De novo* transformer modeling improves recovery of genetic cell types from sparse single-cell RNA sequencing"

Sensitivity to alignment order

We tested ssSTICI's sensitivity to the order of sites in the input CV alignment across all three datasets. We created a random permutation of CV alignment positions by randomizing the column order in the input CV alignment (while retaining the same random seed for determining the order in which rows are processed) and preserving the base calls observed for each cell at each position. We used ssSTICI to predict missing base calls, then restored the original ordering before evaluation.

Model training time was unaffected for the simulated dataset, which fit entirely within one context window and thus precluded convolution. However, for the OC dataset, the model initially converged in fewer than 100 epochs, but after permutation it required over 700 epochs to converge. The permuted TNBC dataset never fully converged even after 1,000 epochs (versus ~80 epochs initially).

The simulated dataset showed very marginal changes (average *NI* decreased by 2.1%, *DR* by 17.7%), while for the OC dataset, performance dropped relative to the unaltered dataset (average *NI* dropped by 0.5%, average *DR* dropped by 52.4%), and the change was small but mixed for the TNBC dataset (average *NI* increased by 6.0%, *DR* decreased by 18.3%). Thus, destroying the original positional relationships between variants did not collapse imputation performance, as ssSTICI still equaled or outperformed other methods, suggesting that ssSTICI prediction accuracy does not depend entirely on physical adjacency between somatic variants.

Sensitivity to masking pattern in training

To test whether ssSTICI base-call accuracy depended on the pattern of sites masked in training, we masked the same percentage of observed bases but changed the random seed used to select masked sites each time. This created five separate predicted alignments (all calls were retained with no filtering to maximize compatibility between runs), which were then used for phylogeny inference. We calculated *NI* and *DR* in each case. These results show that *NI* and *DR* are stable across runs, supporting the conclusion that while ssSTICI is not fully deterministic, it remains highly stable (**Table S1**).

Threefold cross validation

To test whether ssSTICI GT inference and base-call accuracy were sensitive to unobserved positions in the input alignment, we conducted a threefold cross validation in which all sites in the simulated alignment were divided into three bins randomly, and three new ssSTICI models were trained on the remaining third (one third were masked for each model's training). We then used each model to predict base calls (*TBP* = 0.99) for the entire alignment, including the third of sites not observed during training. We observed prediction accuracies of 96.4%, 97.4%, and 97.3% for each model across all sites, and 96.3%, 97.2%, and 97.2%, for the one third of positions which were masked, respectively. We also inferred cellular phylogenies with Fasttree and calculated *NI* and *DR* for each model's predictions, and observed little variation in these metrics (**Table S2**).

**Supplementary Figures & Tables**

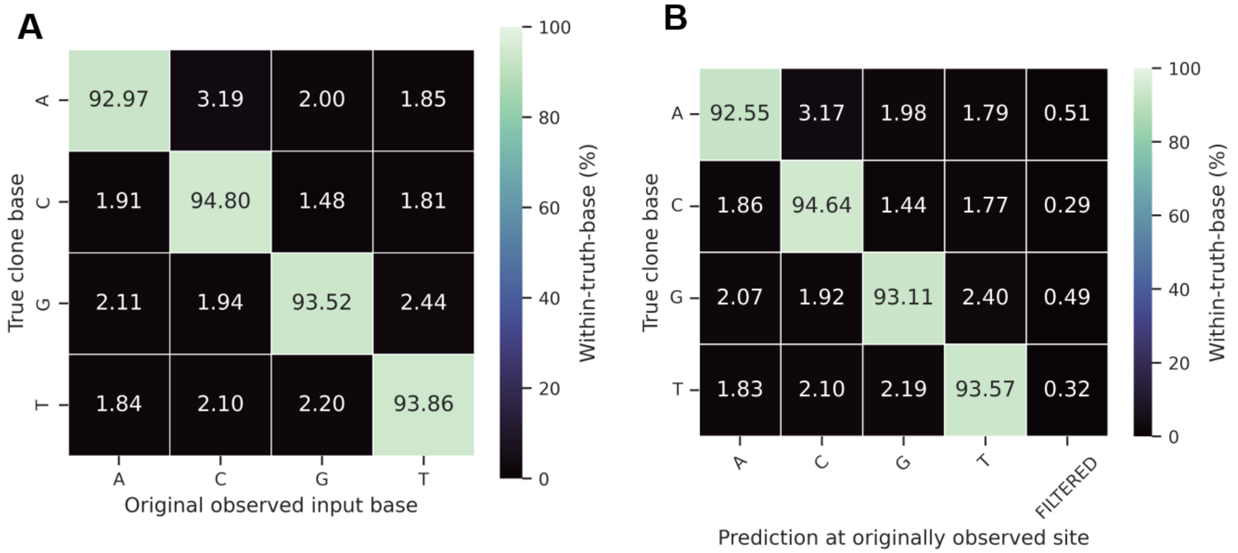


**Figure S1. Confusion matrices for the simulated dataset.**

Frequencies of each base call corresponding to the true base call at that site in the simulated dataset **A)** after we induced error for each true base call in the raw CV alignment and **B)** in the same CV alignment after ssSTICI at a *TBP* of 0.99. The “filtered” column in **B)** counts the percentage sites of with a true base of the given identity which were removed by filtering.

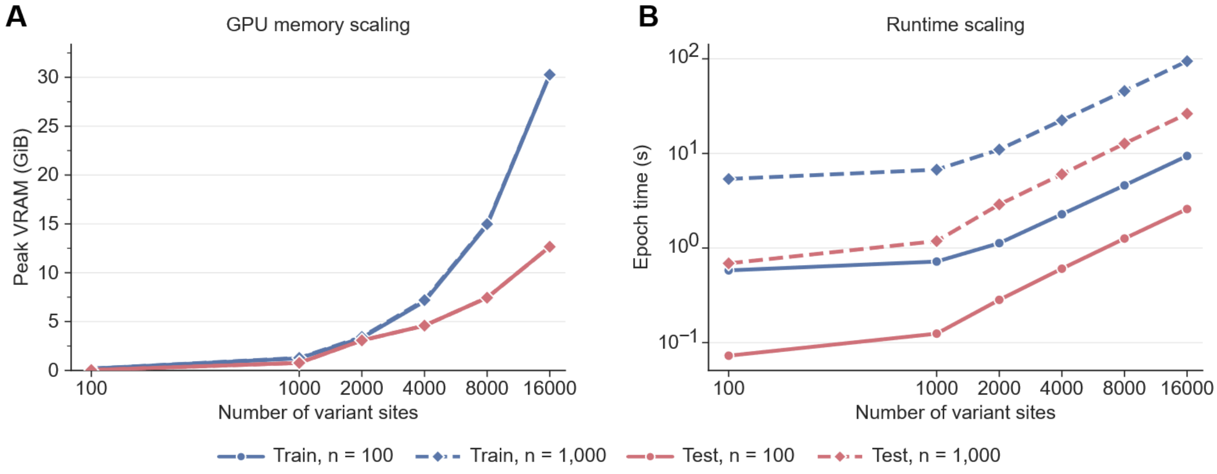


**Figure S2. Resource requirements for ssSTICI.**

Peak VRAM (**A**) scales exponentially with the number of variant sites in the input CV alignment, while the training time per epoch (**B**) scales roughly linearly. We used a batch size of 4 in both cases (*n* = 100, *n* = 1000).

### Supplemental Tables

**A)** *NI*

| GT | A | B | C | D | E | avg | SD |
|---|---|---|---|---|---|---|---|
| 1 | 0.64 | 0.53 | 0.71 | 0.75 | 0.74 | 0.67 | 0.08 |
| 2 | 0.70 | 0.65 | 0.72 | 0.72 | 0.77 | 0.71 | 0.04 |
| 3 | 0.85 | 0.83 | 0.85 | 0.87 | 0.85 | 0.85 | 0.01 |
| 4 | 0.97 | 0.97 | 0.96 | 0.96 | 0.96 | 0.96 | 0.00 |
| 5 | 0.99 | 0.99 | 0.99 | 0.99 | 0.99 | 0.99 | 0.00 |
| avg | 0.83 | 0.79 | 0.84 | 0.86 | 0.86 | | |
| SD | 0.14 | 0.18 | 0.12 | 0.11 | 0.10 | | |

(Columns A–E: Iteration)

**B)** *DR*

| GT | A | B | C | D | E | avg | SD |
|---|---|---|---|---|---|---|---|
| 1 | 6.04 | 4.84 | 4.52 | 4.10 | 5.37 | 4.97 | 0.67 |
| 2 | 2.85 | 2.93 | 2.41 | 2.68 | 2.41 | 2.66 | 0.22 |
| 3 | 3.14 | 3.57 | 3.63 | 3.55 | 2.55 | 3.29 | 0.41 |
| 4 | 7.02 | 5.78 | 5.57 | 6.05 | 4.53 | 5.79 | 0.80 |
| 5 | 7.94 | 9.04 | 7.63 | 7.23 | 6.36 | 7.64 | 0.88 |
| avg | 5.40 | 5.23 | 4.75 | 4.72 | 4.24 | | |
| SD | 2.05 | 2.14 | 1.78 | 1.67 | 1.55 | | |

(Columns A–E: Iteration)

**Table S1. Stability of ssSTICI across runs.**

We trained ssSTICI on the simulated dataset in five separate iterations, masking a different portion of bases randomly in each case. For each run, we then inferred a tree and calculated *NI* (**A**) and *DR* (**B**). We observe that whole-tree metrics (columns, spanning GTs) and per-GT metrics (rows, spanning iterations) were both highly stable for *NI*, and only marginally less so for *DR*.

**A)** *NI*

| GT | Fold 1 | Fold 2 | Fold 3 | avg | SD |
|---|---|---|---|---|---|
| 1 | 0.65 | 0.66 | 0.64 | 0.65 | 0.01 |
| 2 | 0.70 | 0.73 | 0.72 | 0.72 | 0.01 |
| 3 | 0.77 | 0.77 | 0.78 | 0.77 | 0.00 |
| 4 | 0.91 | 0.92 | 0.91 | 0.91 | 0.01 |
| 5 | 0.96 | 0.97 | 0.98 | 0.97 | 0.01 |
| **avg** | 0.80 | 0.81 | 0.81 | | |
| **SD** | 0.12 | 0.12 | 0.12 | | |

**B)** *DR*

| GT | Fold 1 | Fold 2 | Fold 3 | avg | SD |
|---|---|---|---|---|---|
| 1 | 3.59 | 4.11 | 3.70 | 3.80 | 0.23 |
| 2 | 2.32 | 2.42 | 1.99 | 2.24 | 0.19 |
| 3 | 2.19 | 2.71 | 2.11 | 2.34 | 0.27 |
| 4 | 2.30 | 2.85 | 3.27 | 2.81 | 0.40 |
| 5 | 3.14 | 3.64 | 4.64 | 3.81 | 0.62 |
| **avg** | 2.71 | 3.15 | 3.14 | | |
| **SD** | 0.56 | 0.63 | 1.00 | | |

**Table S2. Threefold crossvalidation**

We performed a threefold crossvalidation in which all positions in the simulated dataset were held out, then the typical training protocol with 80% of observed sites masked, and 20% unmasked was carried out. Thus, three unique ssSTICI models were trained each with no knowledge of one third of the dataset. We then used that model to predict GTs in the remaining third and and calculated *NI* (**A**) and *DR* (**B**). Overall prediction accuracies for each fold model were 96.4%, 97.4%, and 97.3%, while prediction accuracies for just the one third of held out sites were 96.3%, 97.2%, and 97.2%, suggesting that holding out some sites did not reduce each ssSTICI model's ability to correctly predict base calls there, nor was overall GT inference disrupted by the masking procedure.
